# Integration of event experiences to build relational knowledge in the human brain

**DOI:** 10.1101/2022.11.17.516960

**Authors:** Anna Leshinskaya, Mitchell Nguyen, Charan Ranganath

## Abstract

We investigated how the human brain integrates experiences of specific events to build general knowledge about typical event structure. We examined an episodic area important for temporal relations, anterior-lateral entorhinal cortex (alEC), and a semantic area important for action concepts, middle temporal gyrus (MTG). Participants underwent fMRI while watching sequences of novel events and recalling their temporal relations over two sessions one week apart. Across different contexts, the roles of these events could either be consistent, or conflict; integrative representations should weaken in the conflicting condition. We found integrative representations that emerged quickly in alEC, and only subsequently in MTG, which showed a significant change across sessions. Thus, alEC is important in building integrative representations early in learning and may be a gateway to semantic knowledge in MTG. We suggest that both episodic and semantic systems participate in integrating new information when it is aligned with their content specializations.

## INTRODUCTION

We use experiences of specific events to build general knowledge about typical event structure. For example, by integrating across multiple instances of making coffee, throwing a ball, or going to a restaurant, we come to know the typical components of these events and their typical temporal relations. Complementary Learning Systems (CLS) theory ^1,2^ proposes that two distinct systems are involved: an episodic system that rapidly encodes recent individual experiences and a semantic system that gradually comes to represent stable properties aggregated across experiences.

The hippocampus (HC) is a critical part of the episodic system. It rapidly binds together the temporal and spatial aspects of specific experiences ^3–9^ but is less critical for longer ago learned information, particularly if it is aggregated across instances ^10–17^. In contrast, semantic knowledge about long-learned, familiar actions and events, such as recognizing the categories *throwing* or *making coffee*, especially relies on areas in lateral posterior temporal cortex surrounding the middle temporal gyrus (MTG; Bedny, Caramazza, Grossman, Pascual-Leone, & Saxe, 2008; Bedny, Caramazza, Pascual-Leone, & Saxe, 2011; Bedny, Dravida, & Saxe, 2013; Bottini et al., 2020; Kable, Kan, Wilson, Thompson-Schill, & Chatterjee, 2005; Leshinskaya, Wurm, & Caramazza, 2020; Tarhan, Watson, & Buxbaum, 2016; Tranel, Kemmerer, Adolphs, Damasio, & Damasio, 2003; Wurm & Caramazza, 2021) ^2^. Cognitively, event and action concepts rely on an understanding of typical relational structure ^27–34^, which offers the hypothesis that relation learning is critical to their formation and as such, may rely on mechanisms in the episodic system to do so. Yet in between episodic and semantic relational representations is a large empirical gap. Our work has shown that MTG represents novel temporal relations learned a week prior ^35^ but it is not well understood by what mechanisms MTG representations are updated with experience nor how episodic and semantic systems might work together to build new relational knowledge.

Here, we test the possibility that this gap is mediated by the contribution of anterior-lateral entorhinal cortex (alEC) and a process of integrative encoding that potentially takes place both in alEC and in MTG. Entorhinal cortex mediates the major efferent and afferent pathways between HC and cortex ^36,37^ and is composed of anatomically and functionally distinct posterior-medial (pmEC) and anterior-lateral portions, with alEC specialized for encoding temporal and object-related information ^38–41^. Connections between EC and HC are essential for temporal associative learning ^42,43^. These findings motivate our hypothesis that alEC has an important role in building temporal relational semantic memory, perhaps as a gateway to the later emergence of relational semantic representations in MTG. However, little evidence exists regarding the relative roles of EC and MTG in the acquisition or retention of relational knowledge and their possible interactions.

Integrative processing is likely critical for building semantic knowledge. By identifying commonalities across diverse specific experiences, neural systems could build generalizable models that enable recognizing the same event structure across instances: for example, making coffee with different methods, or ordering in restaurants that differ in taste and décor. Prior work in memory integration has shown that the ability to link separately presented stimulus pairs that share a common item, e.g., AB and BC, is critically reliant on an intact HC ^44,45^, is correlated with HC engagement during learning ^46–53^, and can result in integrative representations linking A and C ^53–55^. Similar effects are reported in ventro-medial prefrontal cortex (vmPFC; Barron et al., 2020; Schlichting & Preston, 2015; Tompary & Davachi, 2017; van Kesteren et al., 2013; van Kesteren, Fernández, Norris, & Hermans, 2010; van Kesteren, Rijpkema, Ruiter, & Fernández, 2010). However, we expect that integrative representations about event structure, particularly ones that persist longer in time and integrate over distinctly cued contexts, should be seen in the semantic areas that eventually store familiar actions and event concepts (MTG), and perhaps as an intermediary stage in alEC.

While traditional models depict EC as a simple conduit, increasing evidence highlights its role in recurrent processing with HC and in forming more stable and generalizable memory traces that connect diverse experiences ^60–64^. HC might facilitate integration by way of recurrent processing with EC, with EC more likely to hold integrative mnemonic information itself ^43,65,66^. Accordingly, we predict that alEC should also build integrative representations, especially when the learned information must be integrated across distinctly cued contexts and is explicitly about temporal relations, in accordance with its specialization for temporal structure. Such a role for EC would suggest an important modification to the CLS framework and to models of episodic and semantic memory in general.

To test these ideas, we used functional magnetic resonance imaging (fMRI) to measure neural signatures of learning and memory for novel temporal relations across two sessions one week apart. Stimuli were sequences of animated events appearing in the context of one of several novel objects (Figure 1), in which one of the object’s movements (Event A) was reliably followed by one of the other events in the environment (Event B). Participants were encouraged to interpret these temporal relations as the object ‘causing’ the appearance of another event, which our prior work with similar materials has established participants do readily ^67^. We examined learning-related changes during encoding and, subsequently, relational memory strength during recall, as a function of two factors: Session, to test the influence of additional exposure and time, and Consistency, to test for integrative encoding across different contexts or runs. Across different runs, participants saw distinct sequences involving unique objects (Figure 1). Across Consistent sequences, each object movement (Event A) predicted the same outcome Event B, while across Inconsistent sequences, the stimulus serving as Event B varied. If the representation in a neural area is integrative, the memory strength of individual A-B pairs should be influenced by prior A-B pairs, and thus facilitated in Consistent contexts, where they could readily be integrated, while hampered in Inconsistent ones, where they conflicted with prior pairs. This allowed us to investigate the degree of integrative encoding in alEC and MTG both immediately after learning and following more exposure and a week delay.

**Figure 1.**
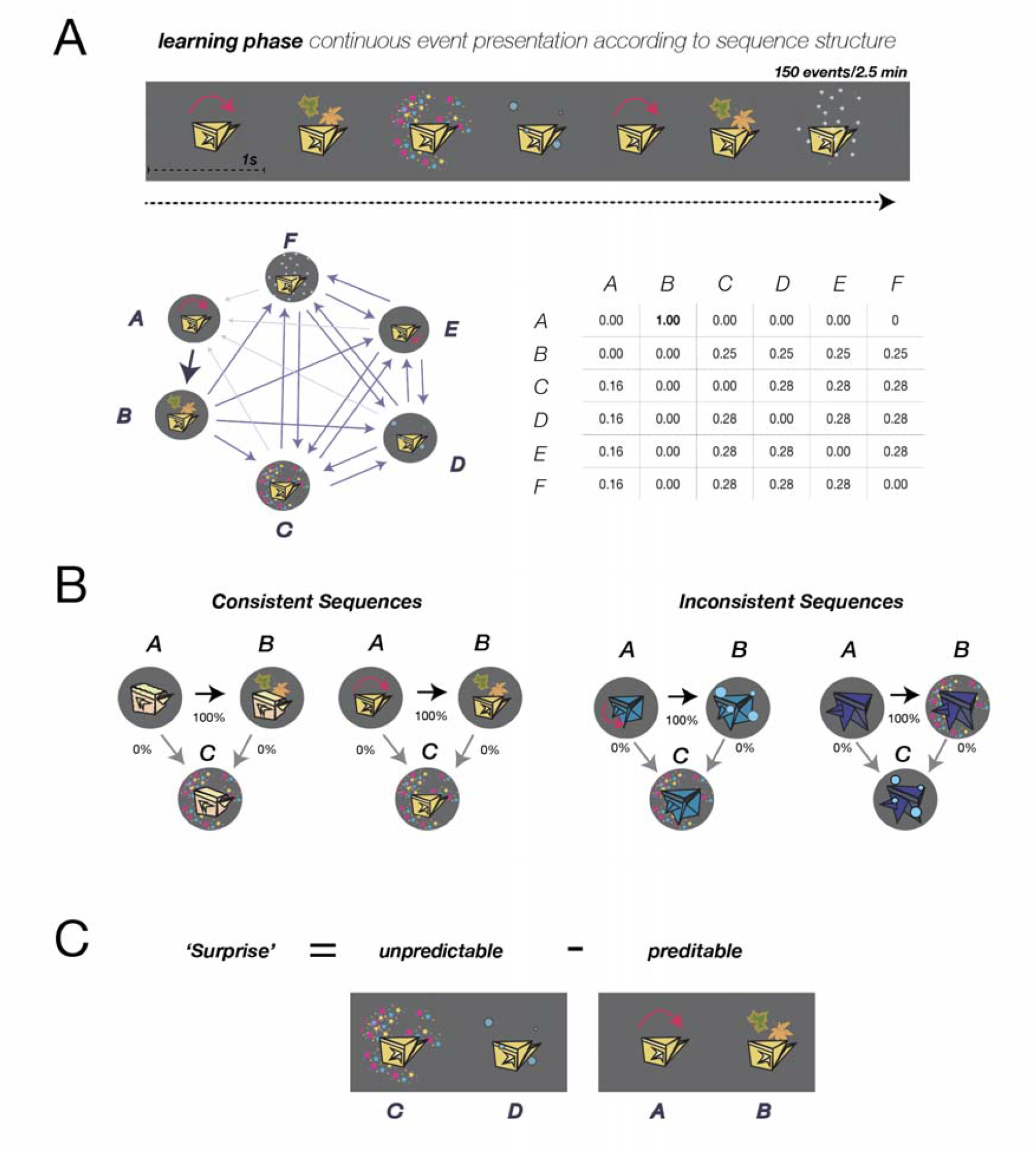
Learning phase materials, task, and analysis design. **A**. Illustration of the Learning Phase, using an example set of stimuli. Participants viewed continuous sequences of animated events that followed a specific transition structure, depicted below as both a graph and a matrix. Event A (always an object movement) was always followed by Event B, but never by other events. Event B, C, and D were ambient (stars, bubbles, snowflakes, etc.), and E was a different object movement. Participants’ task was to identify which ambient event was ‘caused’ by the object’s movement. **B**. Each run of the task showed a unique sequence with a distinct object. Among Consistent sequences, the same ambient stimuli served as Events B and C. Among Inconsistent sequences, Events B and C swapped stimuli, such that if an object movement preceded bubbles in one, it might precede stars in another, creating a conflict. **C**. During the Learning Phase, we measured ‘Surprise’ as a differential response to unpredictable stimuli minus predictable (A-B) stimuli. ‘Change in surprise’ measured how much Surprise strengthened over the course of the learning block.

## RESULTS

### Overview of Paradigm

Participants underwent two sessions of MRI scanning, one week apart, with identical materials. Each run began with a Learning phase (Figure 1A), in which participants explicitly learned temporal relations among a set of events in which Event A (an object movement) was always followed by Event B (an ambient background event such as leaves falling or bubbles floating), but not by Events C or D, which appeared unpredictably. Learning was immediately followed by a Probe phase (Figure 2A), where the same Events (A, B, C and D) again appeared but no longer according to the learned sequence, and instead such that all transitions were equally likely. Thus, any predictive information was only available in memory. Each run pertained to a different sequence with a differently shaped object and belonged either to the Consistent or Inconsistent condition based on its similarity to preceding sequences (Figure 1B). Across Consistent sequences, the relational structure was similar, such that Event B was always the same (e.g., leaves falling). Across Inconsistent sequences, a different stimulus served as Event B (swapping with Event C of the prior sequence), creating a conflict. If the representation of an A-B relation in a given sequence was influenced by prior sequences, and thus integrative, it should be strengthened in the Consistent condition and weakened in the Inconsistent condition. We thus examined the effect of Consistency on neural relational memory strength for individual A-B pairs in the Probe phase and the overall amount of learning-related change in the Learning phase, in Session 1 and Session 2, in several critical ROIs (Figure 3).

**Figure 2.**
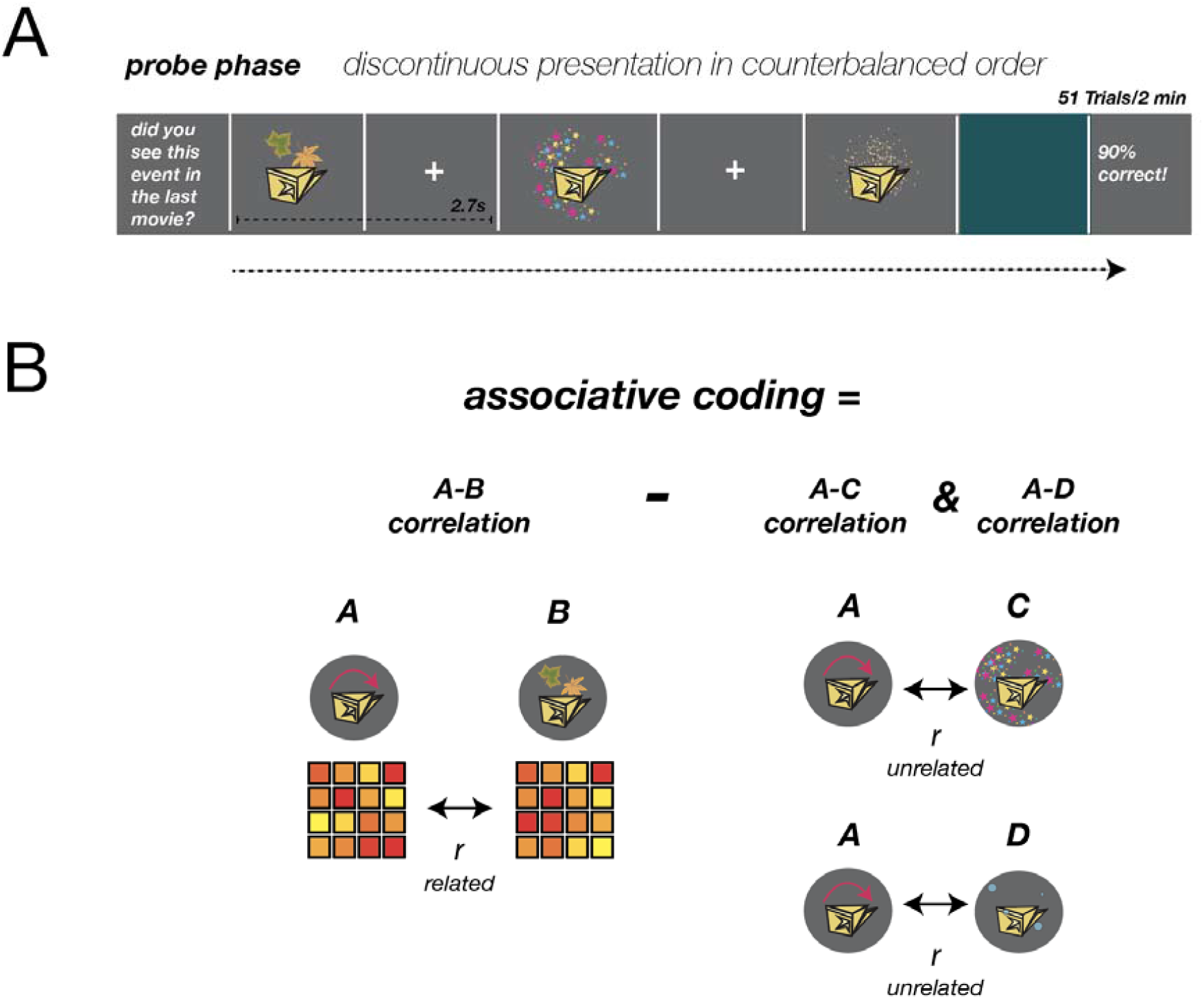
Illustration of the Probe phase task and analyses. **A**. Participants performed a cover task in which they indicated whether each event was one they had seen in the prior Learning phase. The order of events did not follow the learned sequence structure but was rather counterbalanced such that each transition was equally likely, allowing us to estimate the neural response to each event independently of the others. **B**. During the Probe phase, Associative coding was used as an index of relational memory strength between A & B. The neural response to each individual event (A – D) was estimated at each voxel (depicted as the yellow – red squares). In a given ROI, the correlation among the voxel response patterns for each pair of events was computed. The difference in correlation between pairs A & B vs A & C and A & D served as the measure of associative coding.

**Figure 3.**
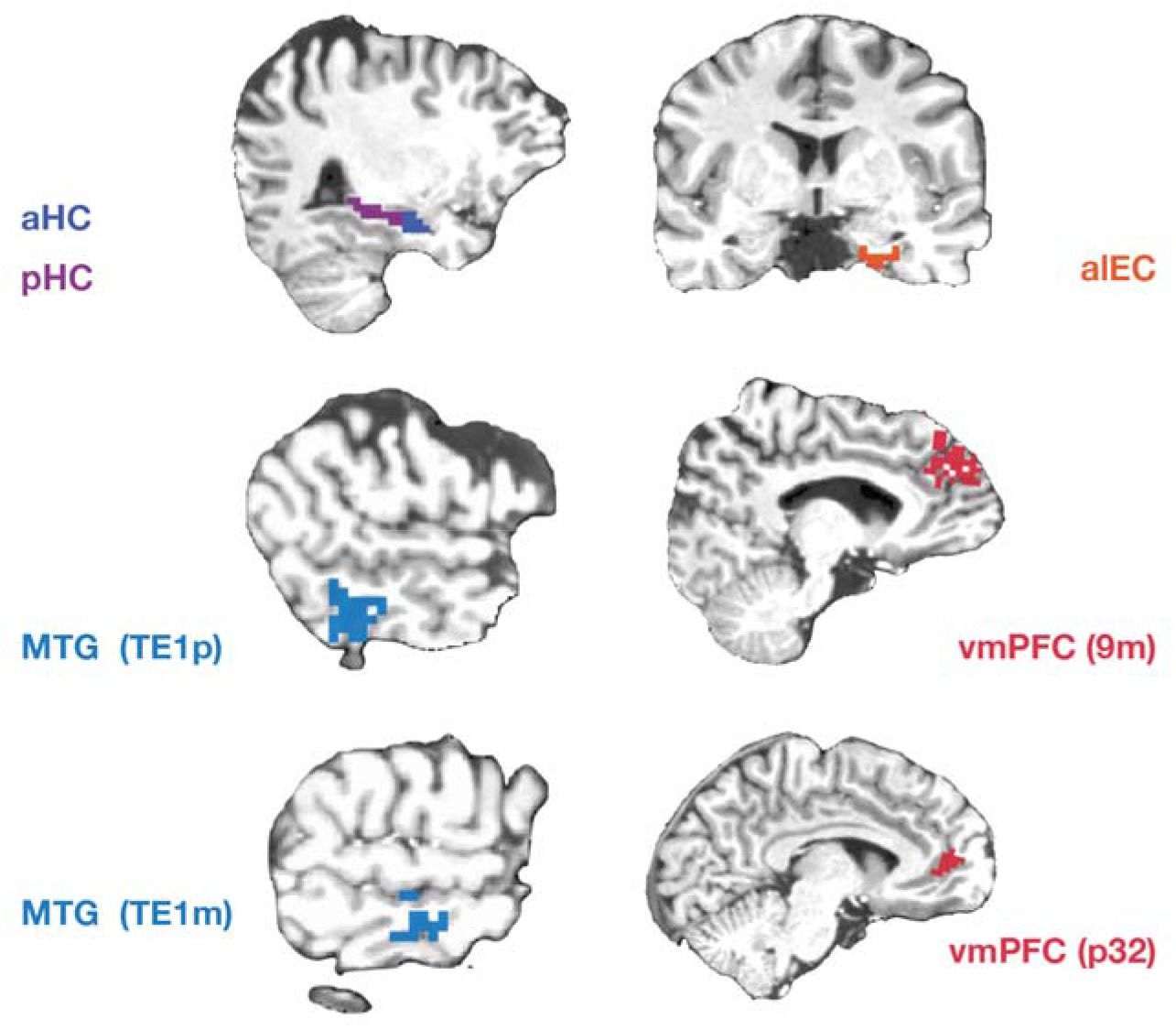
ROIs: hand-traced anterior hippocampus (aHC), posterior hippocampus (pHC), antero-lateral entorhinal cortex (alEC), examples in an individual participant. Middle temporal gyrus (MTG) and vmPFC areas were defined using the surface-based Glasser anatomical atlas ^68^ using areas TE1p, TE1m, 9m, and p32.

These included anatomically defined right-lateralized anterior and posterior HC (aHC, pHC), anterior-lateral and posterior-medial EC (alEC and pmEC), two parts of MTG (Glasser atlas areas TE1m and TE1p, Glasser et al., 2016) and two parts of vmPFC (Glasser atlas areas 9m and p32). Except for vmPFC, ROI definitions were pre-registered, as were analysis methods unless otherwise indicated.

### Behavior During the Probe Phase

During the Probe phase, participants identified whether events were part of the just-seen sequence or not with high accuracy both in Session 1, Consistent *M* = 99.1%, Inconsistent 99.0 %, and Session 2, Consistent *M* = 99.1%, Inconsistent *M* = 99.7 %, indicating high vigilance on the cover task. Analyses reported in the Supplement established that accuracy and reaction time data were not confounded with the fMRI analyses.

During the Probe phase, the order of events was counterbalanced such that all event transitions were equally likely, but it would be expected that cover task reaction times (RTs) would be facilitated for transitions that had been more likely during learning. We thus examined whether RTs to Event B differed as a function of whether it was preceded by the predictive Event A or by unpredictive Events C or D. A Consistency by Transition Type ANOVA in Session 1 revealed a main effect of Transition Type, *F*(1,23) = 5.948, MSE = 0.0252, *p* = .023, indicating that, counter-intuitively, reaction times to Event B were *slower* when preceded by predictive than unpredictive events. Simple effects revealed that this effect was significant within the Inconsistent condition, (23) = 2.29, *p* = 0.031, but not in the Consistent one (*p* > .20). No effects were seen in Session 2. Overall, this shows reverse behavioral facilitation in the Inconsistent condition in Session 1, with unpredictive Events C/D facilitating reaction times to Event B relative to its predictor, Event A, perhaps reflecting the conflict created in this condition.

### Associative Coding During the Probe Phase

We measured relational memory strength for each individual A-B pair using associative coding: a previously reported effect in which events related in memory exhibit a more correlated neural response than unrelated events ^69,70^. During the Probe phase of each run, we obtained an independent measure of the neural response to each Event Type (A, B, C and D), producing for each one a vector of voxel responses in each ROI. Within run, we correlated the vector of voxel responses to Event A with that of Event B, subtracting from this the average correlation of Event A and Event C and of Event A and Event D (Figure 2B). This correlation difference served as our dependent measure, associative coding, separately for each learned sequence/run. Associative coding greater than 0 indicated evidence of relational memory. We tested whether relational memory strength (the magnitude of associative coding) varied by Consistency condition to probe whether these memory representations were integrative, and by Session, to understand how memory varied as a function of exposure and time.

### Anterior-Lateral Entorhinal Cortex

Associative coding in each condition in alEC is shown in Figure 4. If alEC memory representations are integrative, associative coding should be stronger in the Consistent condition than the Inconsistent condition. We also examined the effect of Session on associative coding and on the magnitude of the Consistency effect (as an interaction), to see how memory representations changed as a function of exposure and time. A Session by Consistency ANOVA revealed that associative coding in alEC was higher in the Consistent than Inconsistent sequences, *F*(23,1) = 8.999, *MSE* = 0.323, *p* = .006, with no effect of Session. We further examined effects within Session. There was an effect of Consistency in Session 1, *M =* 0.158, *t*(23) = 3.214, *p* = 0.004, such that Consistent sequences exhibited significant associative coding, *M =* 0.088, *t*(23) = 2.349, *p* = 0.028, while Inconsistent sequences exhibited significant differentiation, that is, negative associative coding, *M* = −0.071, *t*(23) = −3.306, *p* = 0.003. Differentiation entails that Events A and B were less correlated than Events A and C or D. Within Session 2, however, there was no effect of Consistency. Consistent sequences did not show associative coding or differentiation, although there was significant differentiation in Inconsistent sequences, *M =* −0.059, *t*(23) = −2.463, *p* = 0.022. We saw no evidence of associative coding in any condition or any differences in pmEC.

**Figure 4.**
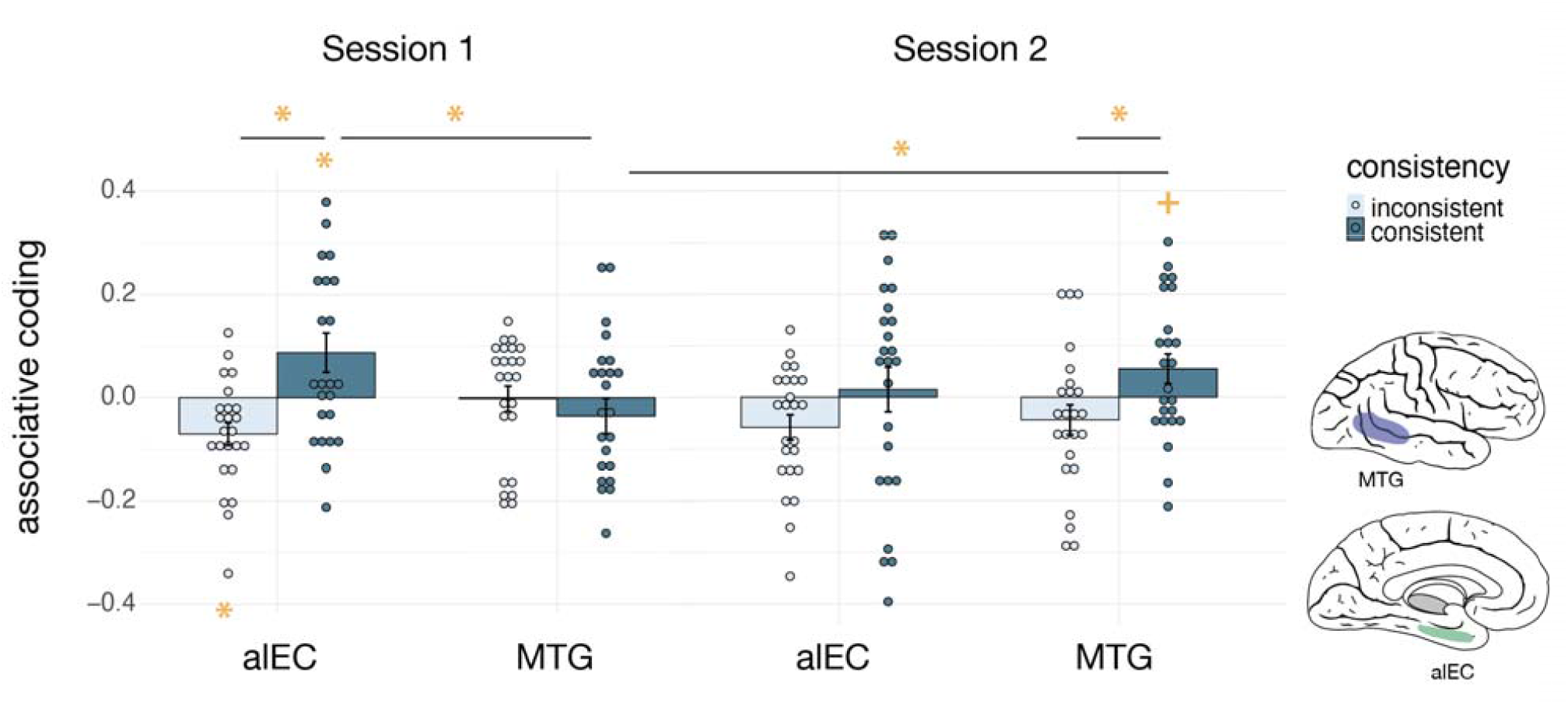
Associative coding for individual A-B pairs in alEC and MTG (TE1p) as a function of Consistency and Session. In Session 1, alEC exhibited associative differentiation in the Inconsistent condition but positive associative coding in the Consistent condition, with a significant difference between these; no effects were seen in Session 2 but there were no interactions with Session. MTG showed significantly less associative coding than alEC in Session 1. In the Consistent condition, associative coding increased in Session 2 vs Session 1 in MTG, yielding a Consistency effect in Session 2. A three-way interaction indicated that these ROIs exhibited effects of Consistency of different magnitudes in Session 1, but not in Session 2. Error bars indicate standard error of the mean (SEM), asterisks denote effects significant at *p* < .05, crosses indicate marginally significant effects.

While surprising, differentiation, in which associated events are pulled apart, is a commonly reported phenomenon ^71–74^ and may especially arise in the Inconsistent condition because relations were conflicting with prior sequences. This neural effect also aligns with the RT findings reported above, which showed inhibition of A-B transitions relative to C/D – B transitions, although there was no correlation across participants for these behavioral and neural measures.

Overall, alEC exhibited associative coding that was highly sensitive to the consistency of information in Session 1, such that it was strongly positive in the Consistent condition and negative in the Inconsistent condition, effects that indicated highly integrative representations. These effects were absent in Session 2 but there was also no significant decline.

### Hippocampus

In aHC, we saw no significant associative coding in any condition (Figure 5). A 2 Session (1 vs 2) by Consistency (Consistent vs Inconsistent) ANOVA did not reveal any effects. There were also no effects of Consistency within Session 1 or Session 2, all *p* > .20.

**Figure 5.**
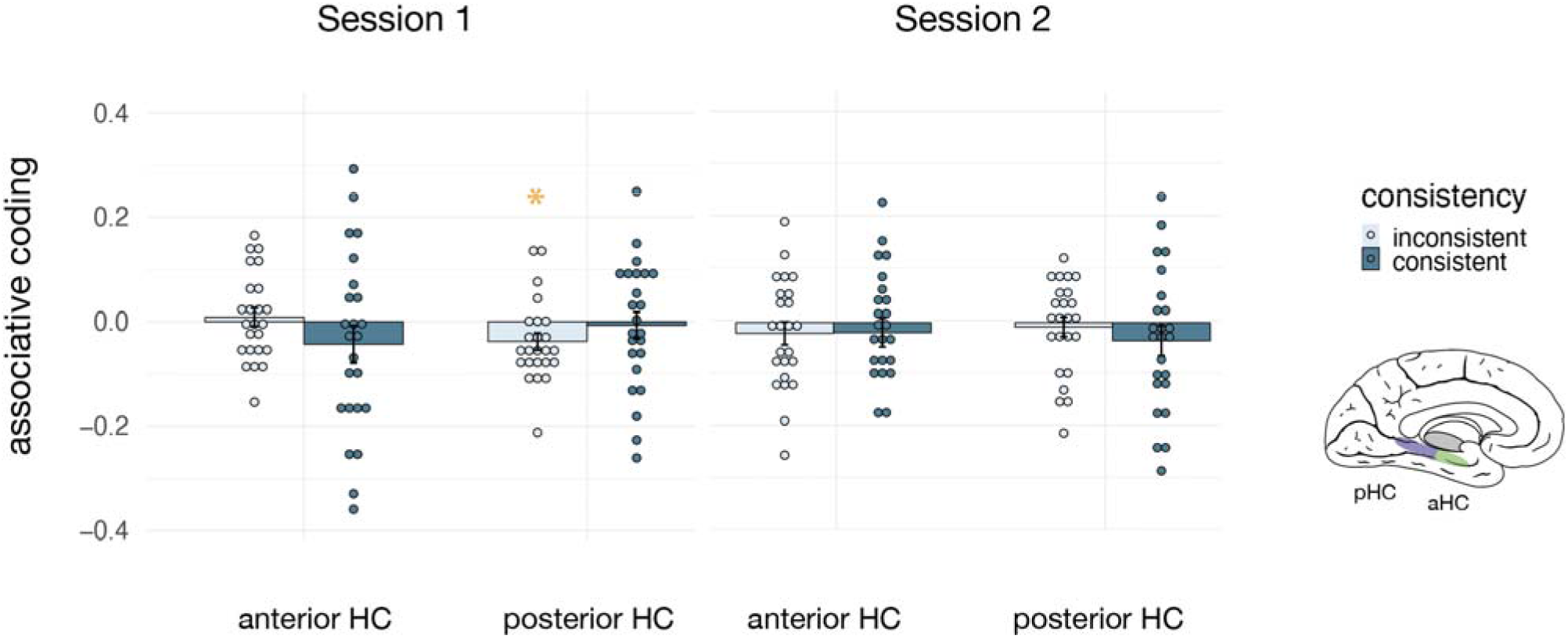
Associative coding (relational memory strength during the Probe phase) in aHC and pHC in Session 1 and Session 2 as a function of Consistency. In Session 1, pHC exhibited negative associative coding (‘differentiation’) such that related events (A-B) exhibited less correlated responses relative to unrelated events (A-C and A-D). No area exhibited positive associative coding. Error bars indicate SEM and asterisks denote effects significant at *p* < .05.

In pHC, a Session by Consistency ANOVA likewise did not reveal any effects and there were no effects of Consistency within Session 1 or Session 2. Associative coding was negative in the Inconsistent condition in Session 1, *M* = −.038, *t*(23) = −2.359, *p* = .027, like in alEC and in alignment with the reported RT findings but also did not correlate with them across individual participants. In summary, there was little evidence of positive associative coding in HC, and no reliable influence of Session or Consistency, indicating little participation of HC in the Probe phase.

Although HC and EC are anatomically interconnected, effects in alEC and pHC appeared divergent. To test this directly, we compared associative coding in alEC and pHC using an ROI by Session by Consistency ANOVA. This revealed a marginal main effect of Consistency, *F*(23,1) = 4.250, *MSE* = 0.167, *p* = .05 and an ROI by Consistency interaction, *F*(23,1) = 8.998, *MSE* = 0.156, *p* = .006, indicating that alEC was more sensitive to Consistency than pHC. This was significant within Session 1, *F*(32,1) = 6.25, *p* = .02, and within Session 2, *F*(23,1) = 4.69, *p* = .041. When alEC was compared with aHC, there was similarly an ROI by Consistency interaction, F(23,1) = 13.99, *p* = .001, which specifically held in Session 1 *F*(23,1) = 13.11, *p* = .001 but not in Session 2, *p* > .10. This reveals overall that integrative encoding was generally higher in alEC than HC.

### Middle Temporal Gyrus

In MTG, using the TE1p ROI, a Consistency by Session ANOVA showed a Consistency by Session interaction, *F*(23,1) = 6.589, *MSE* = 0.106, *p* = .017 (Figure 4). Within Session 1, there was no evidence of associative coding in any condition, nor any Consistency effects, indicating little participation of this area. In Session 2, however, associative coding was significantly higher for Consistent than Inconsistent sequences, *t*(23) = 2.99, *p* = .007. Associative coding in Session 2 was marginal for Consistent sequences, *M* = 0.055, *t*(23) = 1.970, *p* = .061, but not significant in the Inconsistent ones, *M* = −.044, *p >*. 13. Correspondingly, there was stronger associative coding in Session 2 than Session 1 within Consistent sequences, *t*(23) = −2.491, *p* = .020, but not within Inconsistent ones, *p* > .25, explaining the Consistency by Session interaction. This indicates that MTG showed highly integrative representations, but only in Session 2, with a significant increase from Session 1.

We did not see any effects in TE1m, which we had pre-registered but had noted was less likely to show effects based on a prior pilot sample. This suggests our effects are relatively anatomically specific to the more posterior ROI, which is also in line with many of the findings on action and event knowledge in posterior aspects of MTG ^25^. Effects were also specific to the right hemisphere, which we had pre-registered as the focus due to the right-lateralization of prior findings ^35^.

The MTG (TE1p) effect appears qualitatively different from that of alEC. We thus tested whether MTG showed a quantitatively different pattern of effects than alEC using an ROI by Session by Consistency ANOVA over associative coding in the two ROIs. This revealed a main effect of Consistency, *F*(23,1) = 7.388, *MSE* = .266, *p* = .012, an ROI by Consistency interaction, *F*(23,1) = 5.709, *MSE* = .083, *p* = .026, and a 3-way interaction between ROI, Session, and Consistency *F*(23,1) = 6.59, *MSE* = .141, *p* = .017. Within Session 1, there was an ROI by Consistency interaction, *F*(23,1) = 14.39, *MSE* = .220, *p* < .001, while in Session 2 there was a main effect of Consistency, *F*(23,1) = 7.439, *MSE* = 0.180, *p* = .012, and no interactions. This reveals that both ROIs were sensitive to Consistency in Session 2, but in Session 1, alEC was more sensitive than MTG. Follow-up *t-*test showed that associative coding was stronger in alEC than MTG among Consistent objects in Session 1, *t*(23) = 2.488, *p* = .0206, and that the Consistency effect in Session 1 was stronger in alEC than MTG, *t*(23) = 3.794, *p* < 0.001. Overall, this reveals that MTG showed an effect of consistency primarily in Session 2, while alEC did so at both timepoints, and the 3-way interaction demonstrated that these patterns of effects were reliably different between the ROIs.

### vmPFC

Because of the relevance of vmPFC to theories of memory integration ^59^, we performed similar analyses in two anatomically defined vmPFC ROIs (Glasser areas p32 and 9m). In 9m, a Session by Consistency ANOVA revealed no effects (all *p* > .10); within-Session effects were also unreliable (*p* > .07). In p32, there were no effects overall or within Session (all *p >* .14). Associative coding was not significantly positive or negative in any condition. Thus, there was little evidence of the involvement of vmPFC in the Probe phase.

Prior work on vmPFC, however, has often reported mean activation differences, notably more activation during ‘congruent’ or consistent information ^58,75^, rather than effects on the strength of mnemonic information encoding as examined above. In post-hoc analyses to better correspond to prior results, we thus also examined mean activation during the Probe phase as a function of Consistency. In Session 1, we saw marginally more activation in the Inconsistent than Consistent condition, in 9m *t*(23) = −2.059, *p* = .051, and significantly in p32, *t*(23) = −2.208, *p* = .038. This is opposite in direction to prior reports. No mean activation effects were observed in HC, alEC or MTG (post-hoc analysis).

### Searchlights

To identify any additional areas that might show Consistency or Session effects on associative coding, we used whole-brain searchlights. We did not find any clusters passing significance thresholds, but sub-threshold maps reveal that the strongest areas to show a Consistency by Session interaction were in the vicinity of MTG and precentral sulcus (Figure S2, Supplementary data). This suggests that our MTG findings are relatively anatomically specific. Further searchlight results are shown in the Supplementary Data.

### Learning Phase

During the Learning phase at the start of each run (Figure 1A), participants were either initially exposed (Session 1) or re-exposed (Session 2) to predictive information by watching each intact sequence (the memory of which we examined during the Probe phase.) The Learning phase fMRI data provided an opportunity to examine how the roles of our ROIs in memory compare to their roles in learning. To measure learning-related signals, we used a measure of ‘surprise’ (Figure 1C) as the differential activation to expected vs unexpected events, specifically difference between unpredictable events (C or D) and predictable pairs (A-B; because A and B were perfectly auto-correlated, they could not be examined separately). This differential response could be in either direction, where a positive value would reflect a stronger response to unpredictable information and a negative value would reflect a stronger response to familiar information, perhaps due to recollective processes. Either response is expected to scale as participants increasingly learn to identify the predictive pairs. We thus computed ‘change in surprise’ as the difference in surprise between the start and end of each learning phase to measure the amount of learning-related change in each ROI. This was then compared to 0 to measure if learning-related change was reliable, and then compared between Sessions and Consistency conditions. If change in surprise is stronger in the Consistent than Inconsistent condition, then an area exhibited more learning-related change in situations where information could be built up from the preceding sequences than when it conflicted, indicating integrative encoding during learning. Although these measures were pre-registered, we had not pre-registered the comparisons between Consistency conditions or Sessions, so these analyses are exploratory and motivated by comparing them to effects observed in the Probe phase.

### Anterior-Lateral Entorhinal Cortex

Surprise and change in surprise in alEC were negative, indicating a stronger response to predictable than unpredictable events that increased with learning, likely driven by recollective processes engaged during predictive pairs. A Session by Consistency ANOVA on change in surprise revealed a Consistency by Session interaction, *F*(23,1) = 12.100, *MSE* = 0.154, *p* = .002, revealing a Consistency effect within Session 1, *t*(23) = −3.215, *p* = 0.004, but not in Session 2. Session comparisons showed that Consistent sequences showed more change in surprise in Session 1 than Session 2, *t*(23) = −2.256, *p* = .034, whereas Inconsistent sequences showed more change in surprise in Session 2 than in Session 1, *t*(23) = 2.240, *p* = .035, explaining the interaction. In Session 1, Consistent sequences showed increasingly negative surprise, *M* = −0.061, *t*(23) = −3.01, *p* =.006, but Inconsistent sequences showed no change, *M* = 0.037, *p* > .10 (Figure 6).

**Figure 6.**
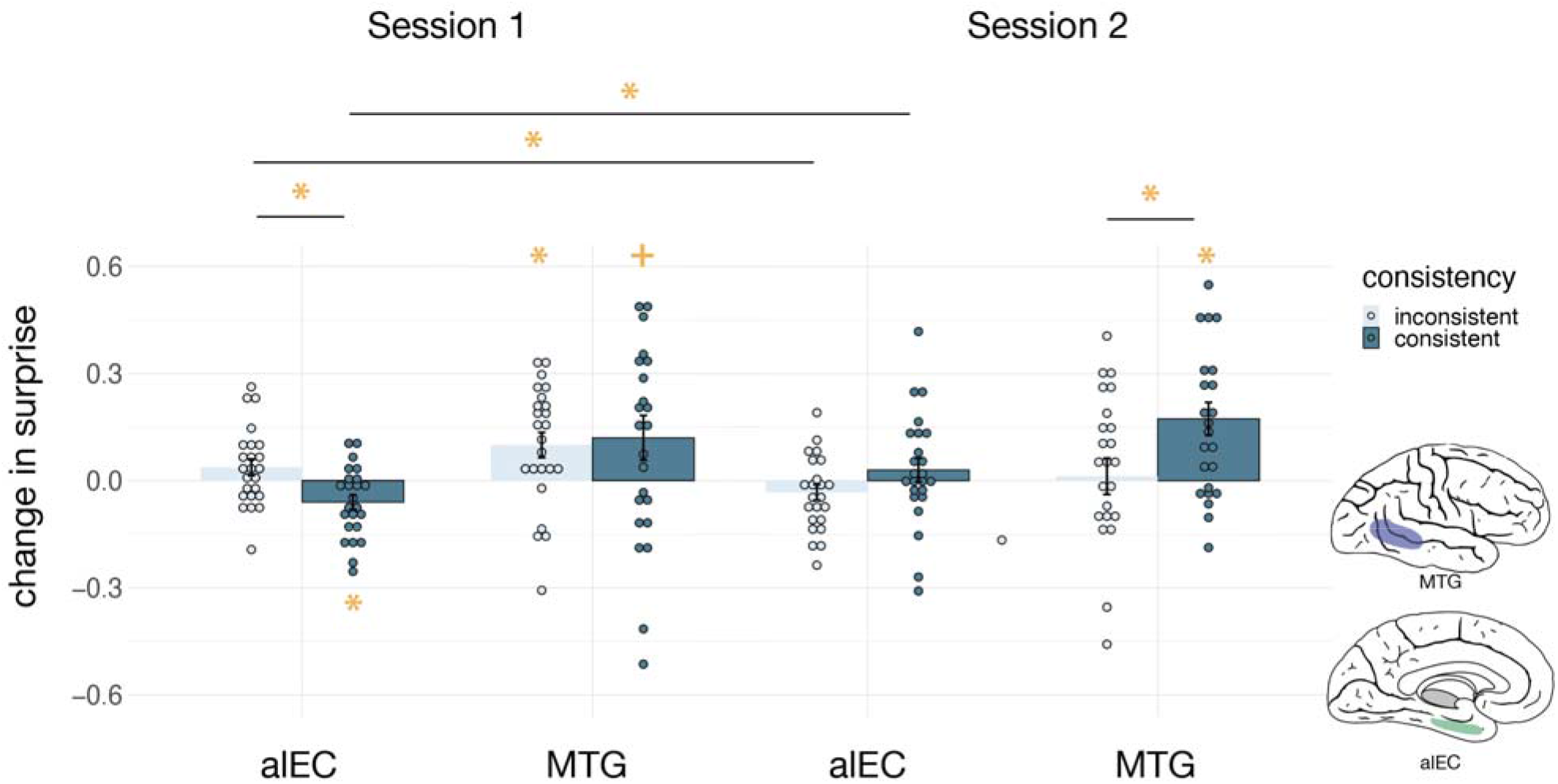
Change in Surprise during the Learning phase in alEC and MTG (TE1p) as a function of Session and Consistency. In Session 1, alEC showed increasingly negative surprise in the Consistent condition, significantly more than the Inconsistent condition, which showed no change. This interacted with Session, such that this difference was absent in Session 2. MTG showed significant change in surprise in the Inconsistent condition in Session 1 with no Consistency difference. A Consistency effect emerged in Session 2, where only the Consistent conditions exhibited significant change. Error bars indicate SEM, asterisks denote effects significant at *p* < .05, and crosses indicate marginal effects.

Change in surprise was not significant in Session 2 in either condition, *p* > .10. Thus, change in surprise in alEC was sensitive to Consistency and Session, with stronger (more negative) effects of Consistency in Session 1 than in in Session 2. This indicates integrative representations that were updated more in Session 1 than in Session 2.

### Hippocampus

aHC exhibited negative change in surprise in Session 1 in Consistent sequences, *M =* −0.057, *t*(23) = −2.535, *p* = 0.019, and marginally so in Inconsistent sequences, *M* = −0.040, *t*(23) = −2.01, *p* = 0.056, but no change in surprise in Session 2, *p* > .20. A Session by Consistency ANOVA did not reveal any effects of Consistency or Session on change in surprise magnitude (Figure 7). Thus, although aHC showed some learning-related changes in Session 1, these did not vary by Session or Consistency. In pHC, there was no significant change in surprise and no effects in a Session by Consistency ANOVA. The observation of negative surprise in HC is consistent with the observation that unpredictability per se does not largely drive HC responses and that instead, recollective processes engaged during predictive trial pairs might dominate the responses here ^76^.

**Figure 7.**
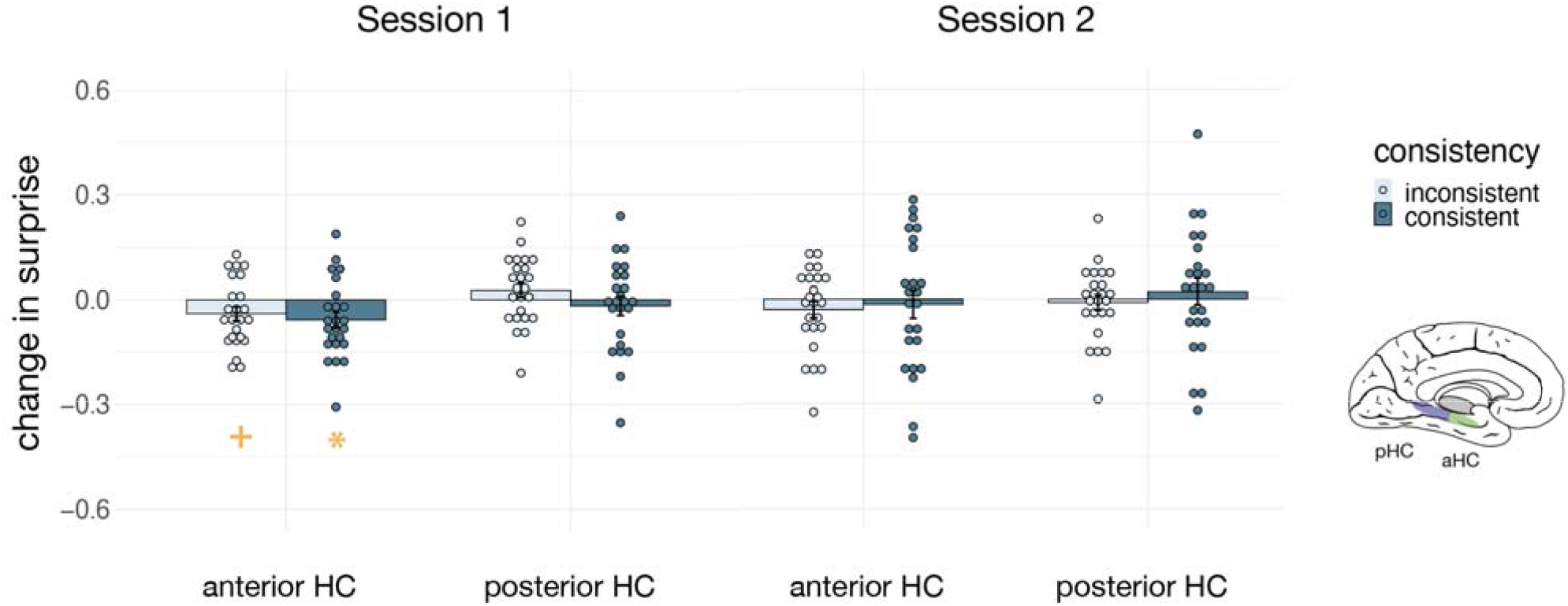
Change in Surprise during the Learning phase in aHC and pHC as a function of Session and Consistncy. Only aHC showed any significant change in surprise, with increasingly negative surprise from the start to the end of the Learning phase in Session 1 in the Consistent condition, but no effect of Consistency or Session and no interactions. Error bars indicate SEM, asterisks denote effects significant at *p* < .05, and crosses indicate marginal effects.

However, there was no evidence of change or condition differences.

The above set of results suggest that alEC might be more sensitive to Consistency than aHC. We tested this directly using a 3-way ANOVA over change in surprise, with factors ROI, Session, and Consistency. This revealed a 3-way interaction, *F*(23,1) = 4.434, *MSE* = 0.049, *p* = .046, and a two-way interaction between Session and Consistency, *F*(23,1) = 6.685, *MSE* = 0.112 *p* = .017. Simple effects revealed that across both ROIs, the effect of Consistency was overall larger (more negative) in Session 1 vs Session 2, *t*(23) = −2.586, *p* = .017. Moreover, the Consistency effect changed more between Session 1 and Session 2 in alEC than aHC, *t*(23) = −2.106, *p* = .046. However, there were not greater Consistency effects overall in alEC than aHC. This suggests these areas did not significantly differ in the extent of Consistency sensitivity but that alEC exhibited more Session differences. This is consistent with the idea that alEC declined in integrative learning by Session 2.

### Middle Temporal Gyrus

Unlike alEC and aHC, MTG exhibited positive change in surprise, i.e., more activation in response to unpredictable than predictable increasing with learning (Figure 6). In Session 1, Consistent sequences showed marginally positive change in surprise, *M* = 0.112, *t*(23) = 1.945, *p* = 0.064, and Inconsistent ones were significantly positive, *M =* 0.100, *t*(23) = 2.900, *p* = 0.008. In Session 2, change in surprise was significantly positive within Consistent sequences, *M* = 0.174, *t*(23) = 3.847, *p* < 0.001, but not within Inconsistent ones, *M* = 0.013, *p* > .80. A Session by Consistency ANOVA on change in surprise revealed only a marginal main effect of Consistency, *F*(23,1) = 3.887, *MSE* = 0.197, *p* = .061 and no interactions. Within Session 1, there was no effect of Consistency, *p* > .70, but there was in Session 2, *t*(23) = 2.511, *p* = 0.020. Overall, MTG exhibited learning-related change activity throughout both Sessions with only some evidence of Consistency differences.

To quantify differences between MTG and alEC, a 3-way ROI by Session by Consistency ANOVA revealed a main effect of ROI, *F*(23,1) = 13.890, *MSE* = 0.559, *p* = .001, an interaction between ROI and Consistency, *F*(23,1) = 6.066, *MSE* = 0.140, *p* = .022, and an interaction between Session and Consistency, *F*(23,1) = 8.642, *MSE* = 0.272, *p* = .007. Within Session 1, there was a main effect of ROI, *F*(23,1) = 7.82, *MSE* = .354, *p* = .01, reflecting that change in surprise was positive in MTG and negative alEC. There was also a trend towards an ROI by Consistency interaction, *F*(23,1) = 3.73, *MSE* = 0.0829, *p* = .066, in the direction of potentially stronger Consistency effect in alEC than MTG. In Session 2, there was a main effect of ROI, *F*(23,1) = 10.08, *MSE* = 0.2138, *p* = .004, again reflecting positive vs negative change in surprise in the two ROIs. Unlike in Session 1, there was a main effect of Consistency, *F*(23, 1) = 6.00, *MSE* = 0.301, *p* = .022, reflecting the fact that both ROIs exhibited Consistency effects in Session 2. Follow-up t-tests revealed that Consistency effects were stronger in Session 2 than Session 1 overall, *t*(23) = −2.940, *p* = 0.007. In summary, MTG showed positive change in surprise while alEC showed negative change, and the ROIs did not differ in the strength of their Consistency effects.

### vmPFC

In vmPFC (9m and p32), there was no significant change in surprise in any condition (*p* > .05) and there were no effects of Consistency or Session (*p* > .10).

To summarize, learning-related changes during sequence exposure revealed effects of Consistency and Session most prominently in alEC, which exhibited learning-related changes more strongly in the Consistent than Inconsistent condition in Session 1, but not so in Session 2, mirroring the memory effects in the Probe phase. MTG exhibited learning-related changes through both Sessions. This aligns with the idea that alEC was not substantially affected by the presentation of learning materials after Session 1, while MTG was, corresponding with its slower updating of memory representations (which became stronger in Session 2). Learning-related changes were minimal in HC and absent in vmPFC, also in line with memory findings.

### Inter-Region Correlations

The findings from the Learning and Probe phases overall suggest that both alEC and MTG played significant roles in learning and memory, but with different time courses, such that alEC showed a rapid formation of memory representations and a later decline in learning, while MTG representations were slower to update and showed continued learning-related change. Our pre-registration included the hypotheses that there would be a functional relationship between alEC in Session 1 and MTG in Session 2, which we tested by examining cross-session correlations of individual participants’ memory strength (associative coding) and learning related change (change in surprise). We saw that change in surprise was negatively correlated between alEC at Session 1 and MTG at Session 2, *r* = −0.565, *t*(22) = −3.209, *p* = 0.004, showing that negative change in surprise in alEC in Session 1 predicted more positive change in surprise in MTG in Session 2. There were no correlations in associative coding (*p* > .70). This suggests some functional relationship between these areas across time, but this did not appear to hold on memory measures, indicating mixed evidence for this hypothesis.

### Post-Scan Recall

Forced-choice questions at the end of Session 2 required participants to recall each of the predictive relations they had learned from all sequences. Questions asked participants to selected between two event pairs: A followed by B vs A followed by C, in the context of the background object cueing the sequence it belonged to. These questions were expected to be difficult because they required participants to recall which relational structure went with which object shape, as event stimuli were highly overlapping across sequences. We found that for pairs from Inconsistent sequences, participants were not overall above chance (50%), *M* = 46%, *p* > .35, but they were significantly above chance for pairs from Consistent sequences, *M* = 56%, *t*(23) = 2.106, *p* = 0.046, with no significant difference between conditions.

In a post-hoc exploratory analysis, we tested which neural signatures might have predicted participants’ ability to retrieve these event associations from their context cues using correlations between individual participants’ accuracy on this test and their neural measures (associative coding and change in surprise) in our ROIs. We found a correlation with associative coding in aHC in Session 1, *r* = 0.467, *t*(22) = 2.476, *p* = .021 and with change in surprise in pHC in Session 2, *r* = −0.444, *t*(22) = −2.323, *p* = .029. In vmPFC, we saw correlations in Session 1 in p32, *r* = 0.435, *t*(22) = 2.2632, *p* = .034 and marginally so in 9m, *r* = 0.389, *t*(22) = 1.983, *p* = .060; effects held across Sessions, 9m, *r* = 0.486, *t*(22) = 2.607, *p* = .016, and p32, *r* = 0.425, *t*(22) = 2.204, *p* = .038, but not in Session 2 alone. These correlations tentatively suggest that HC and vmPFC did have functional relevance to context-cued relational memory in our task, but should be interpreted with caution given their post-hoc nature. No correlations were seen with either learning or memory measures in MTG or alEC.

## DISCUSSION

Using fMRI, we observed that alEC and MTG play complementary roles in the acquisition of relational knowledge. Signatures of relational memory and of integrative encoding were seen immediately after learning in alEC and were followed after more exposure and time by similar effects in MTG, revealing different time courses for similar functions in these areas. Congruently, learning-related change declined with time in alEC but persisted in MTG. These findings offer the new insight that alEC and MTG have important roles in building new, integrative representations of event relations, shedding light on the neural pathways that build semantic memory from experience. This serves to bridge a major gap between episodic encoding, memory integration, and the formation of semantic memory, and aligns with the idea of different specializations within these systems.

### Implications for theories of neural organization of episodic and semantic memory

CLS theory has long proposed that experiences are first encoded in an episodic memory system in MTL (HC and EC) but eventually come to rely on cortical sites elsewhere. Yet prior work has rarely tracked memory representations both in MTL and in specific semantic areas, nor investigated when integrated memory representations emerge in semantic sites. We reasoned that, with exposure and time, encoding of new relational knowledge should emerge in semantic sites specialized for the content of what is learned—here, temporal relation knowledge in MTG, selected on the basis of prior patient and imaging evidence on its role in action and event concepts (discussed further below). We also predicted that MTG should integrate cross-context information. Finally, we probed whether EC might serve as an intermediate stage between episodic and semantic memory. Our findings support these ideas by showing signatures of integrative encoding in MTG that increased with time and exposure, following rapidly formed integrative memory in alEC. The participation of these particular areas aligns with their specializations in temporal episodic memory and action and event semantics, respectively.

alEC exhibited a unique role in rapidly encoding and integrating newly learned temporal relations. It showed evidence of relational memory in the first session, where it also showed a significant effect of consistency, indicating integrative encoding; this was significantly stronger than in HC. These effects are likely related: by virtue of integrating across contexts, it could build stronger representations in the Consistent condition than it could it if it did not integrate. In Session 2, we saw no significant decline in these effects, suggesting alEC’s role may persist across a week’s delay. Correspondingly, learning-related changes were stronger in Session 1 than Session 2, suggesting less updating during the second exposure, and were also stronger in the Consistent condition in Session 1.

MTG showed evidence of relational memory and integrative encoding subsequently to EC, after more exposure and a week’s delay, with a significant increase from Session 1 to Session 2. Learning related changes were seen in both sessions, in line with slower learning. We thus supported the hypothesis that, following sufficient exposure and time, new, integrated relational knowledge increases its reliance on this specific semantic site.

The joint participation of alEC and MTG in these processes supports the idea that the specialized role of alEC in temporal relational memory ^38–41^ is functionally related to the specialization of the semantic site that is eventually updated with similar learning; in this case, MTG to event and action concepts ^25,26,77^, which have relational structure as a core property ^27–29,33,78^. We thus suggest that alEC and MTG together form a potentially specialized system for building event knowledge from experience.

It remains unclear whether the representations in alEC and MTG emerged independently or interactively. Our findings were mixed: we did not find that individual differences in relational memory strength in alEC in Session 1 predicted those in MTG in Session 2, but we did see that Session 1 learning-related changes in alEC predicted those in Session 2 in MTG. Recent theories suggest that new information consistent with prior knowledge is cortically represented more quickly and may update in parallel with HC ^75,79,80^, whereas the classic view is that HC ‘teaches’ cortex ^1,81^. It thus remains an important question for future research to understand when new semantic knowledge in MTG forms independently of alEC and in what ways they might interact.

In contrast to findings from associative inference paradigms ^59,82^, we did not find evidence of integrative representations in HC as measuring during memory or learning, nor much evidence of relational memory or learning-related change. It is possible that weak signatures of relational memory might have arisen because HC did not capitalize on cross-context consistency. Nonetheless, some prior work has in contrast reported integrative memory representations in HC ^53–55^. In these studies, participants learned separate stimulus pairs A-B and B-C; when shown separately during recall, A and C elicited correlated neural responses. Our paradigm differed from these in the way we probed integration and in that we taught separate pairs in distinctly cued contexts: we measured whether associative coding for event pair A-B was stronger when preceded by A’-B than A’-C, when these pairs were shown with diverse background objects. HC is known to elicit diverse responses to similar events when they are associated with unique contextual details ^15,83–85^. Thus, distinct contexts might had led HC to separate rather than integrate different A-B pairs by binding them to their unique contexts. This idea motivated an exploratory analysis in which we correlated individual differences in memory strength in HC with the ability to recall and distinguish all of the A-B pairs at the end of the study based on each context cue. We found tentative, post-hoc evidence for this idea. Another important caveat however is that the lower spatial resolution of our functional data precluded a more detailed examination of HC subfields, among which CA1 is known to be more integrative than others ^86,87^. Future work using high resolution imaging could potentially better resolve conflicting findings regarding integrative coding in HC.

vmPFC is also a common area targeted in studies of memory integration across neurophysiology ^10,75,88–92^ and fMRI ^49,53,55,93^. Again in contrast to prior work, we did not find significant evidence for integrative representations in vmPFC, perhaps for similar reasons as for HC. We saw instead that overall activation was higher in the Inconsistent than the Consistent condition, possibly in line with findings that mPFC is critical for incorporating conflicting information into existing knowledge structures ^89^ and the view that vmPFC supports the process of integration more than serving as a memory site for the integrated content per se ^58,88,94,95^. Nonetheless, it is also possible that vmPFC supports integrated memory representations for specific kinds of content, unlike those here, which were designed to engage MTG.

### Relationship to prior work on MTG

We focused on right-lateralized MTG following our prior work showing relational memory for novel visual events, learned one week prior, in this area ^35^ as well as the many findings connecting this anatomical region to semantic memory for actions, tools, and events ^25,26^. For visual stimuli, MTG responses are often bilateral ^20,96,97^ and include signatures of generalization for action categories across actors ^98,99^, effectors ^100^ or physical manners of execution ^101,102^ and of event memory ^103^. Although these effects surround MTG, it is best considered a mosaic of regions with various specializations. This includes effects of retrieval of action properties of objects ^19,97,104–106^ and, on the left side, selectivity for verbs over nouns and sensitivity to grammatical structure ^20,77,107,108^ as well as selectivity to tools and hands ^109–115^. Our prior work has also shown that temporal relational information is explicitly reflected in tool-selective parts of MTG ^116^. The present findings are within anatomical range of these prior results, but it is not possible to establish if they pertain to the same functional area without a within-study comparison.

### Relationship to prior work on EC

Our work is in line with increasing evidence demonstrating the role of EC in integrative relational memory. Relational memory in general has long been attributed to HC ^4,8^ with EC implicated alongside it by virtue of EC’s role as the major source of afferent and efferent connections between HC and neocortex ^36,37^. Yet recent evidence characterizes EC as more than just a relay, instead playing an active, integrative role in information processing ^43,60,65,66^ and serving as an early bio-marker of Alzheimer’s disease ^117,118^. EC-HC recurrent connections are themselves essential for temporal associative learning ^42,43^ and it is these recurrent connections that may allow EC, more than HC, to form integrative relational memory ^65^. Our findings bolster this view.

Our findings also fit an emerging picture of MTL as composed of areas with distinct specializations, including within EC. Associative coding, the measure used here in which temporally-associated stimuli elicit a correlated neural response, is a classic result in the neurophysiology of EC and surrounding MTL areas (Higuchi & Miyashita, 1996; Messinger, Squire, Zola, & Albright, 2001; Miyashita, 1993; Naya, Yoshida, & Miyashita, 2001, 2003), with convergent findings in human fMRI ^70,124^. The specific roles among these MTL areas have not been well established, but recent evidence has shown remarkable specializations. Evidence from rodent models has supported the idea that pmEC encodes information about spatial context, whereas alEC encodes information about temporal context ^40^, and fMRI data in humans suggests that alEC activity correlates with the precision of temporal memory ^41^. Accordingly, memory representations in alEC reflect temporal proximity among stimuli whereas those in pmEC reflect spatial proximity, when spatial and temporal relations are orthogonalized in learning ^39^. Our findings of associative coding for temporal relations in alEC specifically align with this view, and were the motivation for targeting it distinctly from pmEC.

Our findings also contribute to an understanding of EC’s role in forming generalizable relational knowledge that spans diverse experiences. Observations from navigation studies show that pmEC grid cell representations of space ^125^ have more stable, persistent patterns of firing across diverse environments than HC place cells, allowing pmEC to encode common spatial structures across contexts with diverse sensory details; these and other data support the idea that generalizability is a general property of EC coding for space and beyond ^61^. Integrative encoding is likewise a property of other kinds of EC representations, for example in allowing inferential short-cuts across separately learned but connected relations among social stimuli ^93,126^. Our findings that alEC integrates predictive information across contexts with diverse sensory detail to build integrative knowledge aligns with this emerging understanding of the general properties of EC function.

## Conclusion

The present study sheds light on the neural pathways that build knowledge of temporal event relations from experience. We showed that new temporal relational information is rapidly integrated in alEC, prior to similar signatures becoming detectable in MTG with additional exposure and a week’s delay. This suggests that new experiences lead to integrated memory representations first in an intermediary stage in alEC and then emerge in a specific semantic site—here, a region previously established as important for action and event concepts. These results shed light on specific sites within episodic and semantic memory systems for building temporal relational knowledge and their time- and exposure-dependent changes. We anticipate these findings to advance neural and computational models of memory updating and interaction among episodic and semantic memory systems.

## Acknowledgements

This work was funded by grants ONR N00014-17-1-2961 to C.R. and NSF 2022685 to A.L. We thank Mateo Pitkin for assistance with data analysis.

## Author Contributions

Conceptualization and Methodology, A.L. and C.R; Software, A.L. and M.N.; Formal Analysis, A.L. and M.N.; Investigation, A.L. and M.N.; Resources, A.L. and C.R.; Data Curation, A.L. and M.N.; Writing—Original Draft, A.L.; Writing – Review & Editing, A.L and C.R.; Visualization, A.L.; Supervision, C.R. and A.L.; Project Administration, A.L.; Funding Acquisition, A.L. and C.R.

## Data and code availability

Code for experiments and analysis, as well as processed MRI and behavioral data, is available at the OSF repository at https://osf.io/5xpza/, DOI:10.17605/OSF.IO/5XPZA. Raw data are available on request.

## METHODS

### Participants

30 participants were recruited from the University of California, Davis community and provided written informed consent. Procedures were approved by the UC Davis Institutional Review Board. Twenty-four participants (18 female, 6 male; mean age 24 years) were included in analyses: four were excluded for excessive head motion and two did not complete both sessions. This target sample size was pre-registered. All were neurologically healthy, right-handed, and eligible for fMRI.

### Preregistration

Methods were pre-registered on the Open Science Framework at https://osf.io/5xpza/, DOI:10.17605/OSF.IO/5XPZA. The present report focuses on a subset of the data collected and described in the pre-registration. Deviations and exploratory (additional) analyses are indicated throughout.

### Stimuli & Procedure

Participants took part in two sessions, one week apart, and performed six runs of fMRI scanning in each session (as well as other tasks not reported here). Each run pertained to one of the different sequences and consisted of a Learning phase followed immediately by a Probe phase and lasted 5 minutes.

During the Learning Phase (152 s), participants watched a continuous sequence of 150 1s-long events that was created from six distinct animated stimuli, denoted as Event Types A-F based on their roles in the sequence. Event B was strongly and uniquely predicted Event A, while the appearance of Events C-F was relatively random. To convey this relational structure, the sequential appearance of events was governed by a transition matrix that specified the probability of any event appearing, given the occurrence of any other (Figure 1A). Participants’ task was to identify the predictable event, called “the effect”, which they selected in a 4-alternative forced-choice question at the end of the run from alternatives Events C, D, and F.

Event stimuli were animated gifs. Events A and E were always object movements/state-changes (“object-based” events) and the rest were appearances of elements surrounding the object (“ambient” events). Stimulus assignments to events B, C and D were perfectly counterbalanced such that all analyses controlled for stimulus effects. Events C and D were typically grouped together for analyses given their equivalence in the experiment.

During the Probe phase (140 s), participants’ task was to indicate, for each event, if it had appeared in the just-seen sequence or was novel. Events shown included all the events from the Learning phase except Event E (as it was not of interest to analyses). Additionally, a null event (a turquoise rectangle) was shown as well as Lure events selected randomly from other sequences. Events no longer followed the predictive structure of the Learning phase; they instead appeared in counterbalanced order, such that each event followed every other an equal number of times (exactly 7 for events of interest and 8 times for the null event). There were 51 trials in total. The events also appeared discontinuously: the entire background object disappeared and the animated event was replaced with a fixation cross for 1.7 s with a total ITI of 2.7 s with a .5 s jitter (Figure 2A). This was designed to discourage participants from continuing to learn about any sequence structure among the events.

The sequences (shown in separate runs) differed in the ways the particular stimuli were assigned each Event Type A-F. Sequences 1-3 were Consistent in their relational structure, while Sequences 4-6 were Inconsistent (Figure 2B). Within Consistency conditions, sequences were shown consecutively but the order of the two conditions was counterbalanced across subjects. Thus, participants either saw 1-3 followed by 4-6, or 4-6 followed by 1-3. In either order, Sequence 1 was never ‘consistent’ with anything prior and was thus considered Inconsistent for purposes of analyses, unless indicated otherwise.

The Consistent sequences each used a distinct object-based event as Event A (e.g., tilting, color changing, and rippling). Event B was always the same event (e.g., bubbles), as was event C (e.g., leaves). Event D could vary among the sequences but never conflicted with other event types. The relational structure among events was thus kept consistent, in that events which served the predictable role always stayed the same and those participating in unpredictable roles either continued to do so or were new.

In the Inconsistent sequences, Events A varied exactly as in the Consistent sequences (e.g., were again tilting, color changing, and rippling). However, the stimuli serving the roles of Events B exchanged roles with Events C or D. For example, in Sequence 4, Event B could be stars, while Events C and D were bubbles and leaves. In Sequence 5, Event A would be leaves while events C and D are stars and bubbles, etc. Thus, the relational structure was conflicting among the Inconsistent sequences.

The sequences were distinguished by a unique object present in all of that sequence’s events. To encourage integration, the three sequences belonging to the same Consistency condition were assigned similarly-shaped objects, as depicted in Figure 2. Thus, Sequences 1-3 had three similarly-shaped objects and Sequences 4-6 also had similarly-shaped objects. The specific set of three shapes assigned to each condition were counterbalanced.

Session 2 was almost identical to Session 1, except that it was followed with additional forced-choice questions: participants selected between two snippets of event pairs drawn from one of the six sequences, comparing A-B vs A-C, A-B vs A-D. The correct choice depended on recalling which relations appeared with object shape. Here, Sequence 1 was grouped with the Consistent sequences since at this point, participants had been exposed to all of them and thus Sequence 1 was consistent with the sequences shown afterwards.

### fMRI Acquisition

MRI data were acquired using a Siemens Skyra 3T scanner at UC Davis using a 32-channel coil. Anatomical volumes were acquired with a T1-weighted MPRAGE sequence with 1×1×1 mm voxel resolution, 256 mm field of view, time to repetition (TR) = 1.90 s, and time to echo (TE) = 3.06 ms. Functional data were acquired with a multiband echo-planar imaging (EPI) blood oxygen level-dependent (BOLD) sequence using 64 interleaved slices with a multiband acceleration factor of 2, 3×3×3mm in-plane voxel resolution, 64×64 mm matrix size, TR = 1250s, TE = 24 ms, and flip angle=76 ?. Slices were aligned to −36 degrees from ACPC to minimize anterior temporal distortion. Static fieldmap estimation were performed by collecting 4 volumes in the reverse encoding direction as the main scans.

### Analyses

Preprocessing was performed with the fMRIPrep package with standard defaults as well as freesurfer and AFNI packages. Anatomical scans were skull-stripped and white matter was segmented from gray. Functional data were registered to the pre-processed anatomical scans using the function *flirt*, and head-motion and rotation realignment parameters extracted. Signal outliers were identified and a high-pass filter of 128s was applied.

Functional slices were slice-time corrected and corrected for distortion based on fieldmap estimation. Finally, functional data were smoothed with a 4 mm full-width half-maximum gaussian kernel.

Linear models were used to estimate condition coefficients on fMRI timeseries. Regressors of no-interest included 6 motion and rotation realignment parameters and their first order derivatives; voxels flagged as signal outliers during preprocessing were excluded. Regressors of interest were created for each type of event seen during the Learning phase (A-F) and Probe phase (A, B, C, D, F and lure trials) separately, with null trials and fixation periods as the implicit baseline. Learning phase data were binned by time in order to examine changes during this phase: trials of each event type were assigned to a bin based on their order of appearance, such that bin 1 for event C included the first three appearances of event C, bin 2 the next three, and so on, for a total of 5 bins per event type. Because events A and B were perfectly colinear during the Learning phase, they formed the same regressor, AB.

During the Learning phase, we computed a measure of ‘surprise’ as a contrast between the unpredictable events (C and D) minus the predictable event pair AB, at each time bin. The slope of this measure across time bins were used as a measure of learning, as changes in the magnitude of this difference must be attributable to learning-related processes.

During the Probe phase, we measured relational memory strength using a multivariate measure, associative coding, which compared voxelwise correlations among pairs of events. For each individual condition, the *t*-value of the coefficients from linear modeling for a given event type was extracted in each voxel, reflecting the extent to which that voxel was activated in response to that condition relative to the null trials. For a given region of interest (ROI) this produced a vector of *t*-values for all of the voxels in that ROI. This vector was then correlated pairwise between specific pairs of conditions, here A & B, A & C, and A & D, and then subtracted. The difference in correlation between A&B minus the other two pairs served as the measure of relational memory strength for the A-B pair. This was done in the same fashion for each sequence, performed separately, and then grouped by condition for analyses as follows: Consistent sequences 2 & 3 composed the Consistent condition while Consistent sequence 1 (the first shown), and Inconsistent sequences 4 – 6 composed the Inconsistent condition. For behavioral analyses, Events C and D were combined into a single condition (henceforth “C/D”) as they were functionally identical and served the same role in relevant analyses.

### ROI Definition

The ROIs reported are shown in Figure 3. HC was defined using automatic segmentations performed by Freesurfer, then split into anterior and posterior portions by hand using morphological criteria: the head was labeled as anterior and the body and tail were labeled posterior. This split was motivated by prior observations regarding functional differences between anterior and posterior HC in the memory integration literature and others ^55,86,94,127^. EC was hand-traced, with alEC and pmEC delineated using tracing criteria guided by previous validation studies ^128^ and the split motivated by functional differences between these subregions ^38,39^. Our preregistration indicated that functional differences between alEC and pmEC were expected, with alEC predicted be relevant here given past work showing its role in temporal relational memory ^39^. No effects were seen in pmEC and it was not further considered. For MTG and other cortical areas, we used the Glasser cortical-surface based atlas ^68^ aligned to individual anatomical surfaces to create individual anatomical ROIs. Our pre-registration indicated two Glasser areas for MTG: right TE1p and TE1m, but we noted in our pre-registration that pilot data indicated TE1p to be of particular importance, which held up in these data as well. Our pre-registration also described a functional localizer that did not work and is not reported here. We did not pre-register vmPFC, but chose Glasser ROIs p32 and 9m based on proximity to previously reported results ^52,53^, as this was motivated directly by connecting our work to past findings. The focus on the right hemisphere throughout is based on prior work showing the role of right MTG in associative coding for similar stimuli ^35^. In exploratory analyses, we also examined left-lateralized areas and further address the issue of laterality in the Discussion.

### Searchlight Analysis

Freesurfer software was used to generate inflated cortical surfaces for each participant ^129–131^ which were aligned into a common space and to functional data using AFNI (mapIcosohedron) and algorithms implemented in the Surfing toolbox ^132,133^. Surfing software was also used to define searchlight neighborhoods (curved cylinders that confirm to individual surface topography) of 27 voxels in size. Analyses were then performed in each neighborhood, treated equivalently to an ROI, and results plotted on the cortical surface maps at the center coordinate of each neighborhood.

Multiple comparison correction was performed using permutations over maximal cluster sizes, which tests the probability of obtaining a cluster of a given size by chance alone. Clusters are defined as contiguous activations above a given individual activation threshold (here, *p* < .001). Permutations are created by creating null maps, data that are not expected to reflect real effects, by exchanging condition labels at the linear modeling stage.

However, null maps retain smoothness. Ten null maps were created for each participant, then sampled randomly for group analyses, which were performed as usual. At each of 1000 iterations, a group test is performed and the largest observed cluster size is recorded, creating a null distribution of maximal cluster sizes under the assumption of no meaningful data. Observed clusters can then be evaluated for probability using this distribution.

### Footnotes

2 It is not likely that all of the reviewed effects are in the same functional area, but the broader region can be thought of as a mosaic of highly related functions. See Leshinskaya et al, 2020 for a review.

