## Supplemental Data for "Integration of event experiences to build relational knowledge in the human brain"

SUPPLEMENTARY DATA

*Behavior During the Probe Phase*

Behavior during the Probe phase was analyzed to rule out possible confounds with fMRI data analyses, for example difficulty or attentional vigilance. The fMRI measure during the Probe phase, associative coding, compared the correlation in neural response among different Event Types (A, B, and C/D), and then compared the magnitude of associative coding between Consistency conditions within and between the two Sessions. There were no differences between Event Types, Consistency conditions, or Sessions in accuracy, which was near ceiling.

Reaction times (RTs; log-transformed) were then compared between Event Types and Consistency conditions within and between Sessions, following the fMRI analyses. In Session 1, an Event Type by Consistency ANOVA revealed an effect of Event Type, *F*(2, 46) = 24.27, p < .001, and no effect of Consistency or any interactions. Follow-up t-tests collapsing across Consistency revealed that responses to Event A (*M* = 6.94) were slower than to Event B (*M* = 6.89), *t*(23) = 5.84, *p* < .001 and slower than Events C/D (*M* = 6.90), *t*(23) = 4.675, *p* < 0.001. In Session 2, there was likewise an effect of Event Type, *F*(2,46) = 19.57, p < .001. Follow-up *t*-tests revealed that again, Event A (*M* = 6.87) was slower than Event B (*M* = 6.81), *t*(23) = 5.725, *p* < 0.001 and slower than Events C/D (*M* = 6.84), *t*(23) = 4.840, *p* < 0.001. An ANOVA with Event Type, Consistency, and Session revealed a main effect of Session, with Session 1 (*M* = 6.929) overall slower than Session 2 (*M* = 6.863), *F*(23,1) = 11.17, MSE = .345, *p* = .003, and a main effect of Event Type, F(2,46) = 31.73, p < .001, but no effect of Consistency or any interactions.

The facilitation of predictable Event B relative to random Events C/D is consistent with prior work showing that predictable events are attentionally facilitated, even outside the context in which they follow their predictors, as here (Barakat, Seitz, & Shams, 2013; Otsuka & Saiki, 2020). However, RT *differences* did not follow the pattern of similarity that was measured with associative coding, which measured the extent to which A and B are more similar relative to C/D. By comparing RT similarity, we observed a marginal trend in the opposite direction, such that RTs were more *different* between A and B than A and C/D in Session 1, *t*(23) = 1.849, *p* = 0.077, and in Session 2, *t*(23) = 2.74, *p* = 0.012, which goes against the predicted pattern of neural similarity and thus avoids a confound with fMRI effects.

Questions immediately following each Probe phase were used to ensure that participants correctly recalled that Event A predicted Event B, not Event C, D, or F, in the sequence they had learned about in that run. Accuracy was near-ceiling in all conditions, Session 1: Consistent *M* = 97.9%, Inconsistent M = 91.7%; Session 2: Consistent 95.8%, Inconsistent, 97.9%. To rule out any accuracy differences between conditions, we performed a Session by Consistency ANOVA, which revealed no effects. To similar ends, we also performed paired *t*-tests within Session and within Consistency, which revealed only that Inconsistent items improved in accuracy between Session 1 and Session 2, *t*(23) = -2.77, *p* = 0.011. This suggests equivalent recall accuracy across Consistency conditions and Sessions.

*Searchlight Findings*

Searchlights for associative coding in each condition, their differences, and interactions were performed to examine relational memory outside of the a priori ROIs. No significant effects were observed for comparisons of Session or Consistency. However, within Consistent sequences, we observed several areas exhibiting negative associative coding or differentiation (Figure S1). These were significant in right precentral, consistent with inhibition between predictively related events relative to unrelated ones (Figure 6). These results are in line with behavioral findings indicating inhibition of A-B transitions relative to C/D-B transitions and prior reports of cortical inhibition of associative links in similar paradigms (Barron et al., 2016; Koolschijn et al., 2019). No other effects passed threshold.

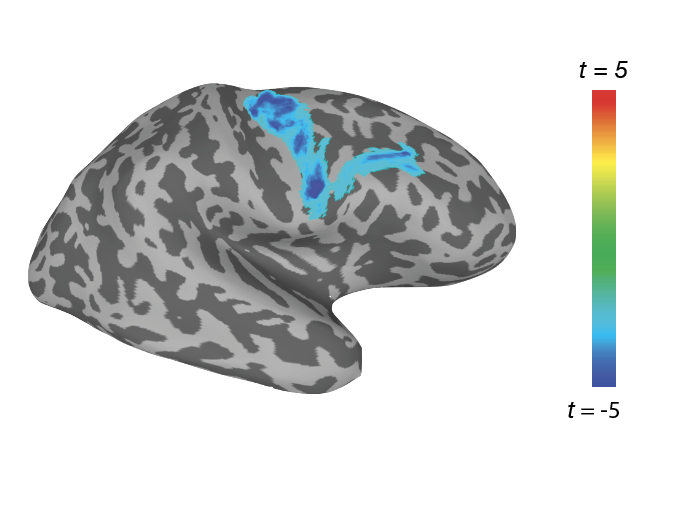

Figure S1. **A.** Whole-brain searchlight results (thresholded at two-tailed cluster-corrected *p* < .05) for associative coding in the Consistent condition at Session 1, revealing *negative* results in right prefrontal cortex.

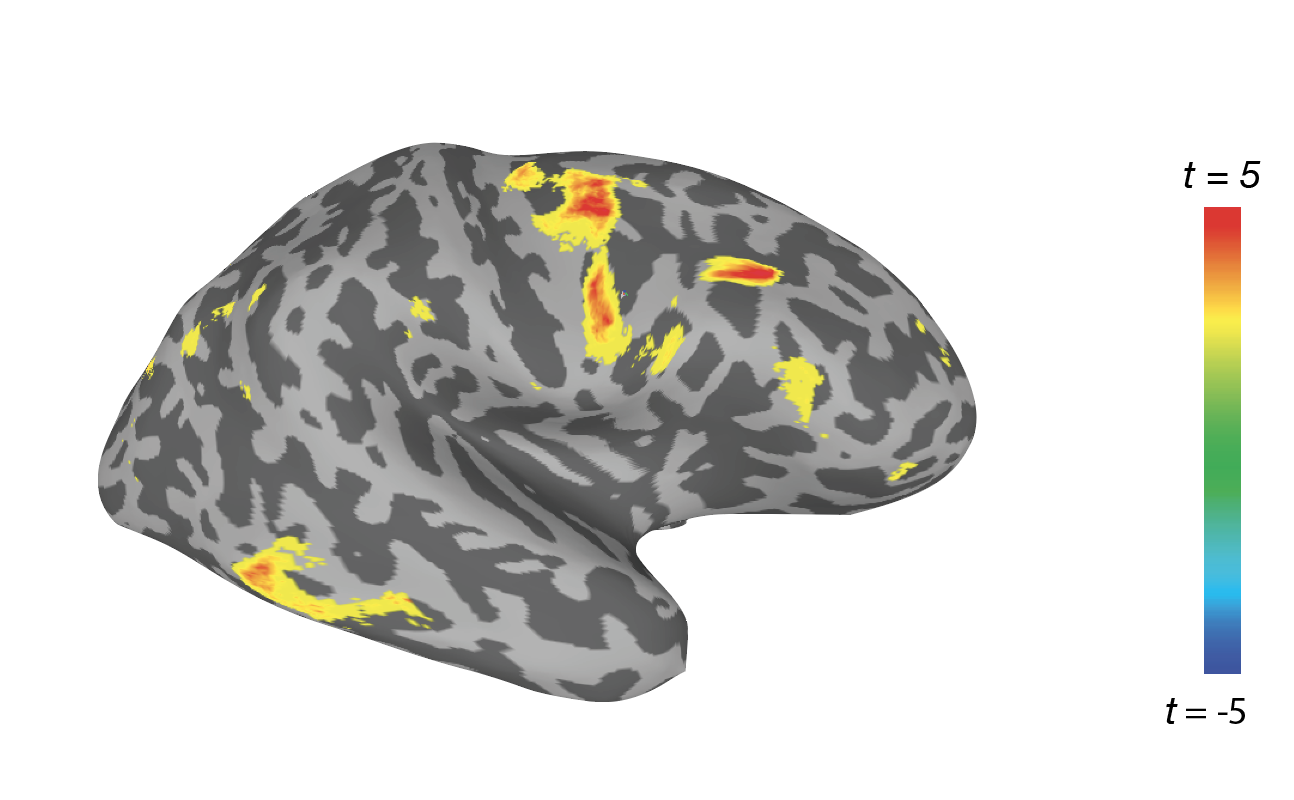

Figure S2. For illustrative purposes only, whole-brain searchlight results for stronger Consistency effects in Session 2 than Session 1, thresholded at an uncorrected level of *p* < .01 and cluster size > 400mm^2^.
